# Selection against expression noise explains the origin of the hourglass pattern of Evo-Devo

**DOI:** 10.1101/700997

**Authors:** Jialin Liu, Michael Frochaux, Vincent Gardeux, Bart Deplancke, Marc Robinson-Rechavi

**Affiliations:** Department of Ecology and Evolution, University of Lausanne, 1015 Lausanne, Switzerland; Swiss Institute of Bioinformatics, 1015 Lausanne, Switzerland; Laboratory of Systems Biology and Genetics, Institute of Bioengineering, School of Life Sciences, Ecole Polytechnique Fédérale de Lausanne (EPFL)

## Abstract

The evolution of embryological development has long been characterized by deep conservation. Both morphological and transcriptomic surveys have proposed a “hourglass” model of Evo-Devo^1,2^. A stage in mid-embryonic development, the phylotypic stage, is highly conserved among species within the same phylum^3–7^. However, the reason for this phylotypic stage is still elusive. Here we hypothesize that the phylotypic stage might be characterized by selection for robustness to noise and environmental perturbations. This could lead to mutational robustness, thus evolutionary conservation of expression and the hourglass pattern. To test this, we quantified expression variability of single embryo transcriptomes throughout fly *Drosophila melanogaster* embryogenesis. We found that indeed expression variability is lower at extended germband, the phylotypic stage. We explain this pattern by stronger histone modification mediated transcriptional noise control at this stage. In addition, we find evidence that histone modifications can also contribute to mutational robustness in regulatory elements. Thus, the robustness to noise does indeed contributes to robustness of gene expression to genetic variations, and to the conserved phylotypic stage.

---

Phenotypes can vary even among isogenic individuals in homogenous environments, suggesting that stochastic effects contribute to phenotypic diversity^8,9^. Gene expression variability among genetically identical individuals under uniform conditions, hereafter “variability”, is one of the most important stochastic processes in the mapping of genotype to phenotype. It is caused by a combination of molecular noise (stochastic biochemical effects, e.g., transcriptional burst process based transcriptional noise) and other effects (variation in cells and their environment, e.g., distribution of molecules at cell division)^10–13^. Precise regulation of gene expression is notably important during development^14^, however, this process inevitably has to deal with stochasticity^15^. This tension between precision and stochasticity in development raises questions, such as whether some stages are more robust to gene expression stochasticity. And whether natural selection against expression variability can transfer to mutational robustness, causing the evolutionary conservation of the phylotypic stage. To answer these questions, we investigated expression variability across fly embryonic development.

We generated 288 single embryo 3’ end transcriptomes using BRB-seq^16^, at eight developmental stages covering the whole fly embryogenesis, with 3h intervals (Figure 1A). After quality control, 239 samples were kept (Figure S1, S2). On average, we obtained over 5 million uniquely mapped reads of protein coding genes per embryo. Based on multidimensional scaling analysis (MDS), 150 embryos follow the developmental trajectory, while there is a small cluster of 89 embryos collected at different time points mixed together (Figure 1B). The samples in this cluster appear to be unfertilized eggs (Methods and Figure S3). All further analysis was performed only on the 150 fertilized embryos.

**Figure 1:**
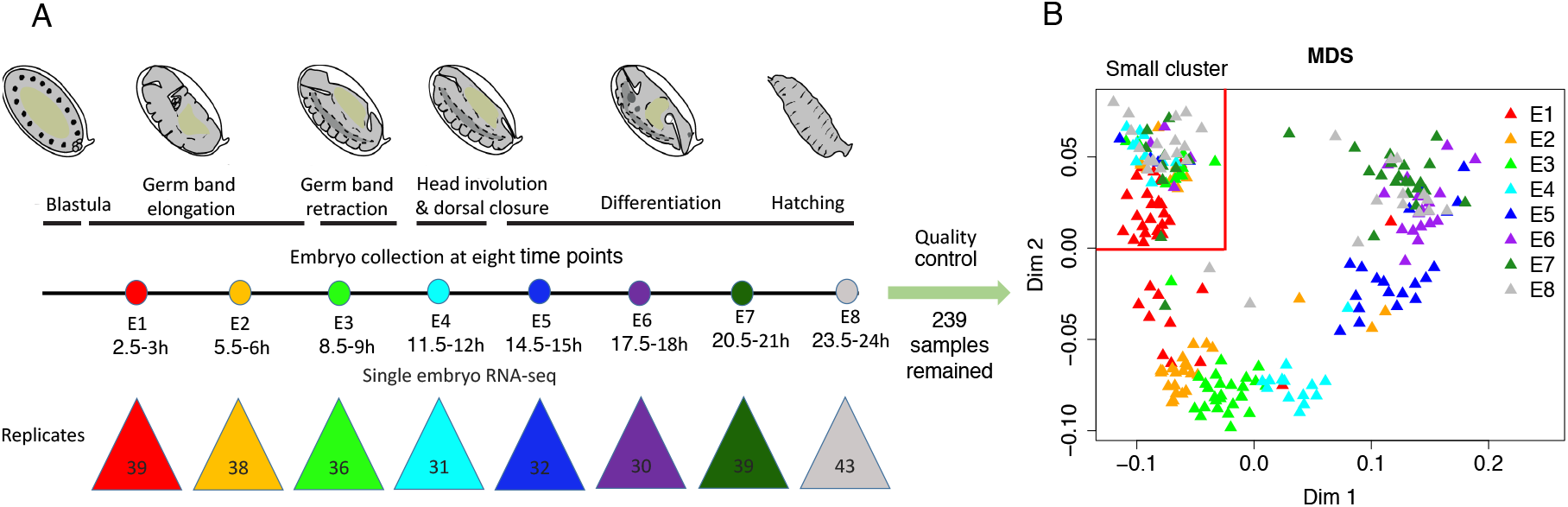
Studying expression variability throughout embryogenesis. A. Method outline. We performed single embryo BRB-seq^16^ at eight developmental stages, indicated by different colored dots. The number of samples collected at each stage is indicated in the colored triangles. Embryo images adapted from^17^ and used with permission from Springer Nature (License Number: 4547630238607) and from the authors. B. Multidimensional scaling analysis (MDS) of 239 high quality samples. Different colors indicate different stages. The samples can be split into two groups: a small cluster in the top-left delimited by two red lines; and the remaining samples, which are organized according to embryonic stage. Only the 150 samples which follow embryonic stages were used for further analysis.

We measured expression variability as Adjusted SD, standard deviation (SD) of expression between replicates corrected for expression level (Methods and Figures S4-6). This expression variability follows an hourglass pattern overall, with a global minimum at E3 (Figure 2A), corresponding to the phylotypic stage of fly^7^. There is also a local minimum at E6. This is consistent with the pattern of transcriptome divergence between fly and mosquito *Anopheles gambiae*, with the global minimum at E3, and a local minimum at E6^18^. Our observations are robust to the use of different variability metrics (Figure S7), and to sampling (bootstrap analysis, Figure S8). Bootstrap results also suggest that the minimum of variability extends over E3 to E4. The embryo transcriptome is dominated by zygotic transcripts 2.5h after egg laying^19^, so the high variability in E1 and E2 is not directly caused by maternal transcripts. We didn’t find any significant functional enrichment for genes which follow the hourglass variability pattern. Overall, expression variability is not equally distributed throughout embryogenesis, and gene expression at the phylotypic stage appears more robust to stochastic factors than at other stages.

**Figure 2:**
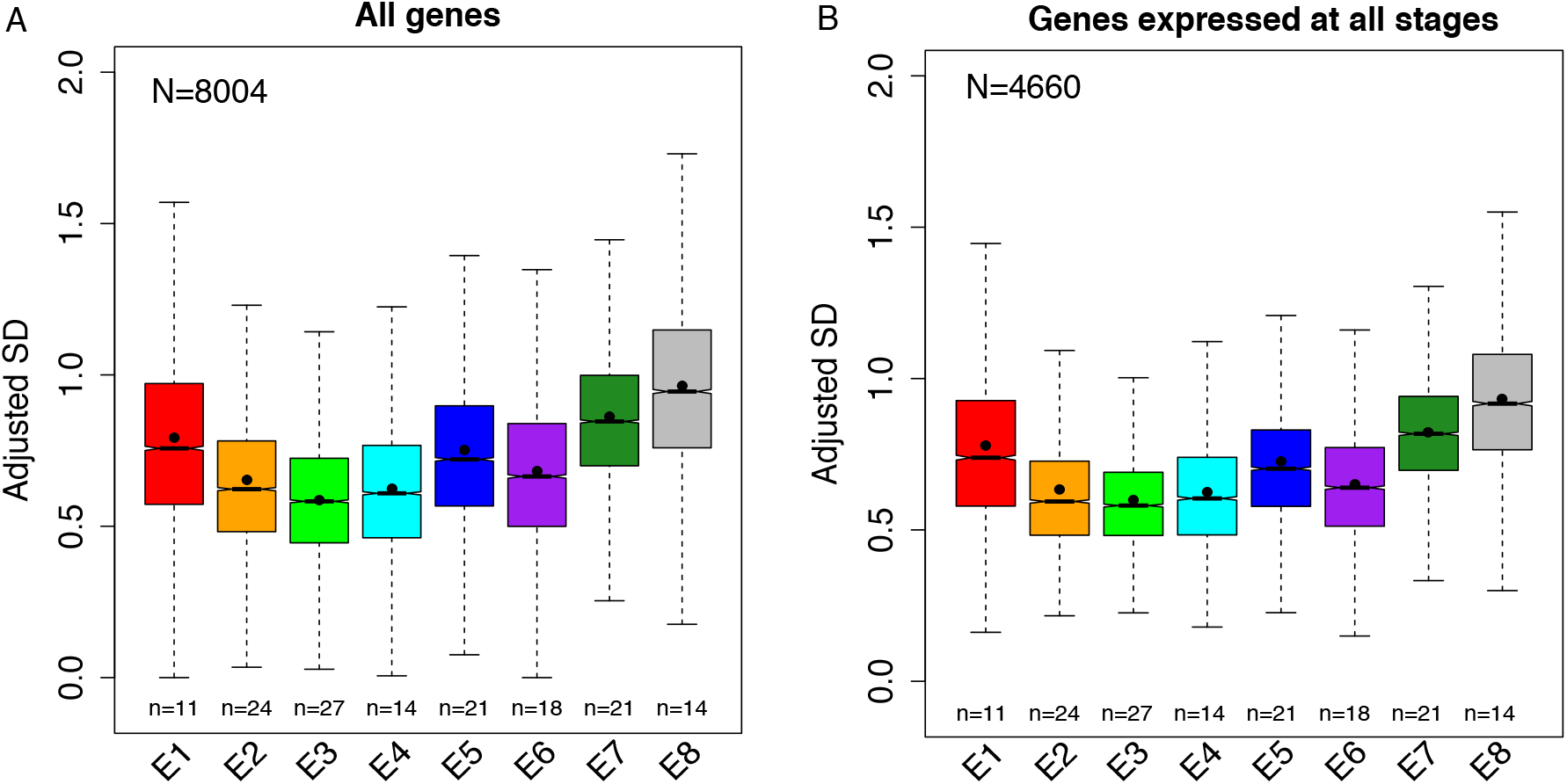
The phylotypic stage (E3) has lower expression variability. The number of individual samples used in each development stage is indicated below each box. The number of genes analyzed is indicated in the top-left corner of each plot. The lower and upper intervals indicated by the dashed lines (“whiskers”) represent 1.5 times the interquartile range (IQR), and the box shows the lower and upper intervals of IQR together with the median. The black dot in each box indicates the mean. A. Expression variability pattern of all genes which passed quality control. We performed pairwise Wilcoxon tests between any two stages to test the significance. The multiple test corrected p-values (Benjamini–Hochberg method) are shown in Table S1; they are all < 10^−7^. B. Expression variability pattern of genes expressed at all stages. We performed pairwise Wilcoxon tests between any two stages to test the significance. The multiple test corrected p-values (Benjamini–Hochberg method) are shown in Table S2; they are all < 10^−5^ except E2 vs. E4, for which p-values = 0.24.

The variation in expression variability could either be due to changes in the set of active genes, with genes differing in their intrinsic variability levels, or to genome-wide changes in the regulation of variability. To test this, we first reproduced our results restricted to the subset of genes which are expressed at all stages. Under the first explanation, we would expect to lose the hourglass variability pattern, but the pattern is maintained (Figure 2B). We performed additional tests: restricting to genes with constant expression level over development (Figure S9A); restricting to transcription factors (Figure S9B); and contrasting genes with dispersed or precise promoters (Figure S10), following Schor et al^20^. Dispersed promoters seem to be more robust to mutations, which might also translate into robustness to noise. Despite a loss of power with fewer genes, there remains an hourglass pattern of expression variability in all cases. Interestingly, the precise promoter genes have higher variability than the dispersed promoter genes except at E3, thus a strongest hourglass pattern. Overall, these results suggest that the lower variability at E3 is due to genome-wide regulation mechanisms more than to changes in the gene set.

Histone modifications can regulate transcriptional noise^21–25^, notably through the modulation of transcriptional burst frequency^22–24^. For example, high levels of histone modifications can increase chromatin accessibility, leading to an increase in transcriptional burst frequency, which leads to minimizing noise. To check this role of histone modifications, we analyzed four available euchromatin histone modifications at six developmental stages^26^. For each gene, we calculated the mean modification signal (background-subtracted tag density) separately for proximal promoters and for gene bodies^23^. Higher modification signal genes tend to have lower variability for all histone modifications (Figure 3A). This supports a role in minimizing transcriptional noise, and is consistent with previous studies in yeast and mammals^22,23^.

**Figure 3:**
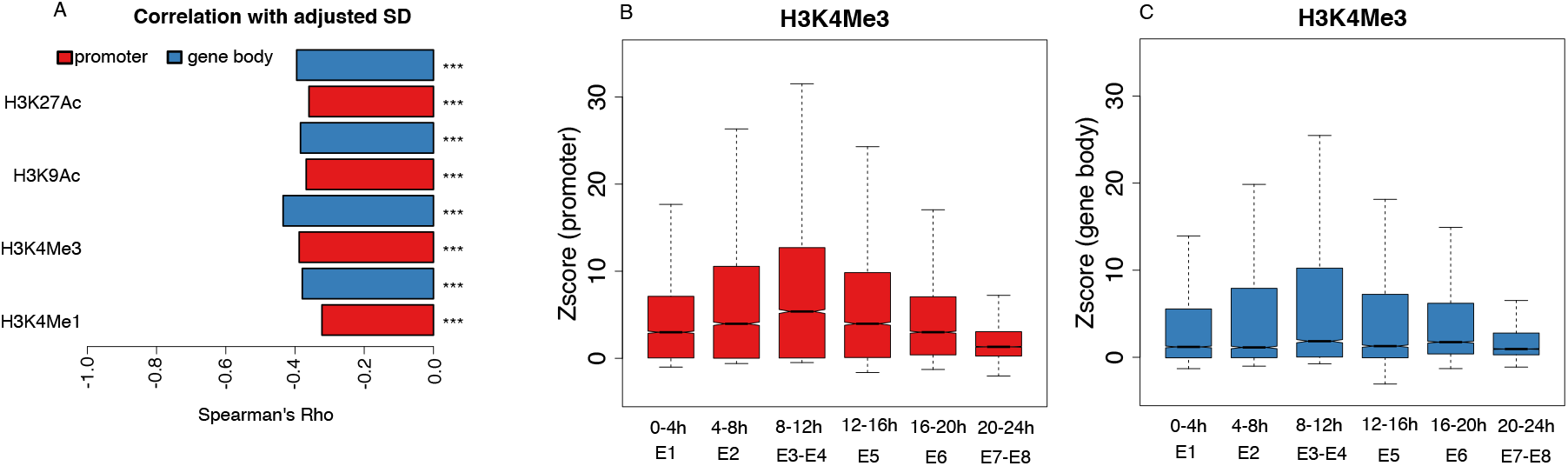
Histone modification signal and expression variability. Red and blue represent histone modification signals calculated on the proximal promoter (4kb around the transcription start site – TSS) and the gene body, respectively. A. Spearman’s correlation coefficient between histone modification signal (background-subtracted tag density) and expression variability. Here, for each gene, both its variability and its histone modification signal are the mean value across stages. ***, P < 0.001; **, P < 0.01; *, P < 0.05, NS, P ≥ 0.05. B. Proximal promoter H3K4Me3 signal (Z score relative to intergenic signal) in different stages. Corresponding stages of our expression variability data are indicated below. The lower and upper intervals indicated by the dashed lines (“whiskers”) represent 1.5 times the interquartile range (IQR), and the box shows the lower and upper intervals of IQR together with the median. We performed pairwise Wilcoxon tests between any two stages to test the significance. The multiple test corrected p-values (Benjamini–Hochberg method) are shown in Table S3; they are all <10^−7^ except 4-8h vs. 12-16h, for which p-value = 0.68. C. Gene body H3K4Me3 signal (Z score relative to intergenic signal) in different stages. Corresponding stages of our single embryo BRB-seq data are indicated below. The lower and upper intervals indicated by the dashed lines (“whiskers”) represent 1.5 times the interquartile range (IQR), and the box shows the lower and upper intervals of IQR together with the median. We performed pairwise Wilcoxon tests between any two stages to test the significance. The multiple test corrected p-values (Benjamini–Hochberg method) are shown in Table S4; they are all < 10^−5^ except 0-4h vs. 20-24h, for which p-value = 0.26; and 8-12h vs. 16-20h, for which p-value = 0.26.

The gene-level relation between histone modifications and expression variability raises the possibility that the pattern of expression variability across development could be driven by changes in histone modification signal. To compare histone modification signal between stages, we normalized gene and promoter signal by that on intergenic regions (Methods), which are not expected to change histone modification signal between stages. All histone marks present an hourglass-like pattern, with the highest signal at 8-12h (except for H3K4Me1 on gene body, where it is a local but not global maximum), corresponding to E3 and E4, i.e. the lowest expression variability, for both promoters and gene body (Figures 3B-C, S11). Moreover, for all histone marks on gene body, as well as H3K4Me1 on promoters, there is another local maximum at 16-20h, corresponding to E6. Generally, these results support changes in histone modification signal over development, with a correspondence between stronger histone modification signal and lower expression variability.

Several studies have suggested that mechanisms which confer robustness to stochastic variation can also buffer the effects of genetic variation^14,27,28^. If histone modifications can buffer the effect of genetic variation on gene expression, we should observe that genes with higher histone modification signal are less sensitive to mutations in their regulatory regions, and are thus less conserved. Indeed, genes with higher histone modification signal tend to have less conserved core promoter sequences^29^ (49 bp upstream TSS and 10 bp downstream from the TSS) between species (phastCons score; Figure 4A). They are also less conserved within a population (promoter nucleotide diversity π; Figure S12). The phastCons pattern remains using 200 bp or 400 bp regions, but disappears using 1 kb regions (Figure S13), indicating a relatively narrow region around the TSS under this balance of selection and robustness.

**Figure 4:**
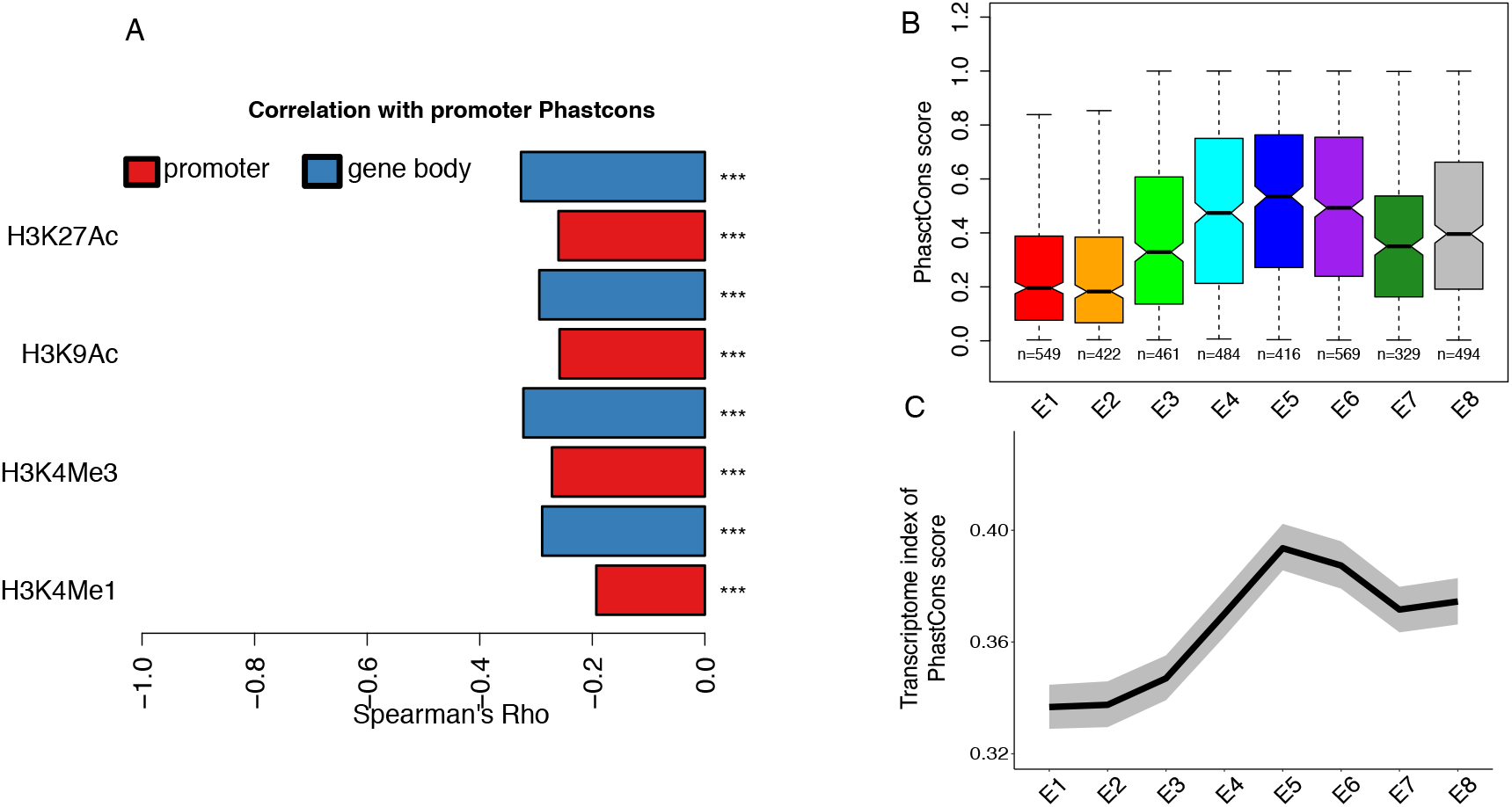
Histone modification signal and promoter sequence conservation. The promoter sequence conservation is the mean of the phastCons score over experimentally identified core promoter regions (49 bp upstream TSS to 10 bp downstream of the TSS)^29^. A. Spearman’s correlation coefficient between histone modification signal (background-subtracted tag density) and promoter sequence conservation. Red and blue represent histone modification signals calculated from the proximal promoter (4 kb around the TSS) and gene body respectively. Here, for each gene, the histone modification signal is the mean value across stages. ***, P < 0.001; **, P < 0.01; *, P < 0.05, NS, P ≥ 0.05. B. Variation of promoter sequence conservation for stage specific genes. The number of genes in each development stage is indicated below each box. We performed pairwise Wilcoxon test between any two stages to test the significance. The multiple test corrected p-values (Benjamini–Hochberg method) are shown in Table S5. C. Transcriptome index of promoter phastCons score across development. The grey area indicates 95% confidence interval estimated from bootstrap analysis.

Since histone modifications appear to buffer genetic variation in gene expression, and since the E3 stage has stronger modification signals, the lower expression divergence in E3 between species^7^ might be caused either by stronger purifying selection on mutations in regulatory regions, or by histone modifications buffering the consequences of mutations in these regions. In the first case, we expect genes specifically expressed at E3 to have higher sequence conservation on promoters. In the second case, we expect the opposite pattern, since mutations that are buffered would behave nearly neutrally. To test this, we identified genes specifically expressed in each stage and compared their promoter sequence conservation. We found that genes specific of E3 have a relatively weak promoter sequence conservation (Figure 4B), supporting a stronger buffering mechanism rather than stronger purifying selection on sequences. The transcriptome indexes of conservation and of diversity (mean promoter sequence conservation and mean π, respectively, weighted by expression) extend this observation to the full transcriptome (Figure 4C; Figure S14). These results support a role of buffering effects on regulatory mutations in the hourglass pattern of expression divergence in fly embryogenesis. Essential genes, and highly connected genes, have lower variability (Figure S17), which supports that variability is detrimental, and that mechanisms which reduce it are adaptive. Thus natural selection on robustness against expression variability could contribute to the phylotypic stage conservation at macroevolutionary scale.

We have found an uneven distribution of variability, and thus of robustness of the process of gene expression, across development, which mirrors the hourglass Evo-Devo model^1,2^. Stage E3 is the most robust to stochastic variation on gene expression, with lower expression variability, and is the phylotypic stage of fly, with conservation between species^7^. Although mutational robustness can evolve under natural selection theoretically^30^, the conditions are too restrictive to be relevant in practice. We propose that the mutational buffering effect of histone modifications is a by-product of selection for minimizing transcriptional noise. Thus, our model is that selection for robustness to noise and environmental perturbations in a key embryonic stage has led to the evolutionary conservation over large time scales which characterizes the phylotypic stage.

## Methods

### Availability of code

Data files and analysis scripts are available on GitHub: https://github.com/ljljolinq1010/expression-noise-across-fly-embryogenesis.

### Availability of data

Expression datasets have been deposited to the Gene Expression Omnibus with accession number <u>GSE128370</u>.

### Embryo collection and RNA extraction

Fly lines (w^1118^) were obtained from the Bloomington stock center and reared at room temperature on a standard fly medium with 12 hours light dark cycle. The fly medium we used is composed of: 6.2 g Agar powder (ACROS N. 400400050), 58.8 g Farigel wheat (Westhove N. FMZH1), 58.8 g yeast (Springaline BA10), 100 mL grape juice; 4.9 mL Propionic acid (Sigma N. P1386), 26.5 mL of Methyl 4-hydroxybenzoate (VWR N. ALFAA14289.0) solution (400 g/L) in 95% ethanol, 1 L Water. 100 to 150 flies were transferred to cages, which were sealed to a grape agar plate (1:1 mixture of 6% agar and grape juice). We used 4 separate cages to collect the embryos. The adults were kept overnight on this plate before being transferred to a new plate supplemented with yeast paste. Synchronization of eggs on this plate lasted for 2 hours before being swapped with a new plate supplemented with yeast paste. We let the adults lay eggs for 30 min before removing the plate and letting the eggs incubate for the desired time. Eggs were harvested using the following protocol. First a 1:1 bleach (Reactolab 99412) 1x PBS mix was poured on the plate and incubated for 2 min. During this incubation, we used a brush to lightly scrape the surface to mobilize the embryos. We then poured the PBS-bleach mixture through a sieve, washed the plate with 1x PBS, and poured the wash on the same sieve. We washed the sieve several time with 1x PBS until the smell of bleach disappeared. Single embryos were then manually transferred to Eppendorf containing 50 *µ*L beads and 350 *µ*L Trizol (lifetechnologies AM9738). The tubes were homogenized in a Precellys 24 Tissue Homogenizer at 6000 rpm for 30 seconds. Samples were then transferred to liquid nitrogen for flash freezing and stored at –80°C. For RNA extraction, tubes were thawed on ice, supplemented with 350 *µ*L of 100% Ethanol before homogenizing again with the same parameters. We then used the Direct-zol™ RNA Miniprep R2056 Kit, with the following modifications: we did not perform DNAse I treatment, we added another 2 min centrifugation into an empty column after the RNA Wash step, finally elution was performed by adding 8 *µ*L of RNAse-free water to the column, incubation at room temperature for 2 min and then centrifugation for 2 min. RNA was transferred to a low-binding 96 well plate and stored at - 80°C.

### Bulk RNA Barcoding and sequencing (BRB-seq)

The BRB-seq is a technique for multiplexed RNA-seq^16^ which is able to provide high-quality 3’ transcriptomic data at a low cost (e.g. 10-fold lower than Illumina Truseq Stranded mRNA-seq). The data (fastq files) generated from BRB-seq are multiplexed and asymmetrical paired reads. Read R1 contains a 6 bp sample barcode, while read R2 contains the fragment sequence to align to the reference genome.

#### 1. Library preparation

RNA quantity was assessed using picogreen (Invitrogen P11496). Samples were then grouped according to their concentration in 96-well plates and diluted to a final concentration determined by the lowest sample concentration on the plate. RNA was then used for gene expression profiling using BRB-seq. In short, the BRB-seq protocol starts with oligo-dT barcoding, without TSO for the first-strand synthesis (reverse transcription), performed on each sample separately. Then all samples are pooled together, after which the second-strand is synthesized using DNA PolII Nick translation. The sequencing library is then prepared using cDNA augmented by an in-house produced Tn5 transposase preloaded with the same adapters (Tn5-B/B), and further enriched by limited-cycle PCR with Illumina compatible adapters. Libraries are then size-selected (200 - 1000 bp), profiled using High Sensitivity NGS Fragment Analysis Kit (Advanced Analytical, #DNF-474), and measured using Qubit dsDNA HS Assay Kit (Invitrogen, #Q32851). In total, we generated four libraries. For details of library information, please check Table S20.

#### 2. Sequencing

Libraries were mixed in equimolar quantities and were then sequenced on an Illumina Hi-Seq 2500 with pair-end protocol (read R2 with 101 bp) at the Lausanne Genomic Technologies Facility.

### RNA-seq analysis

#### 1. Generating expression matrix

The fastq files were first demultiplexed by using the “Demultiplex” tool from BRB-seqTools suite (available at https://github.com/DeplanckeLab/BRB-seqTools). Then, we trimmed the polyA sequences of the demultiplexed files by using the “Trim” tool. Next, the STAR aligner^31^ was used to map the trimmed reads to the reference genome of fly *Drosophila melanogaster* (BDGP6, Ensembl release 91^32^). Finally, the read count of each gene was obtained with HTSeq^33^.

#### 2. Filtering samples and genes

Low-quality samples need to be filtered out, since they might bias results of downstream analyses. In order to assess sample quality, we calculated the number of uniquely mapped reads and of expressed genes for each sample^34^. We removed samples with <0.3 million uniquely mapped reads or with <4500 expressed genes (Figure S1). We confirmed that these filtered samples are indeed outliers in a multidimensional scaling analysis (MDS) (Figure S15). Since lowly expressed genes have larger technical error, to minimize the technical noise, we need to remove lowly expressed genes as well. We first calculated counts per million (cpm) with the edgeR package^35^ for each gene. Then we removed genes with mean cpm across samples ≤1, as suggested by Lun et al.^34^. Finally, for the remaining genes, we re-transformed their cpm values to the original count values for the downstream normalization analysis. After filtering, we obtained an expression count matrix with 239 samples (Figure S2) and 8004 protein coding genes.

#### 3. Normalization and batch effect correction

Because BRB-Seq retains only the 3ʹ end of the transcript, we performed sample normalization by using quantile normalization with log transformation in the voom package^36^, but without transcript length normalization. To remove potential batch effects across the four libraries, we applied the ComBat function in the sva package^37^ to the normalized and log2 transformed expression data. For genes with expression values less than 0 after Combat, or with original expression values equal to 0, we change its values to 0 after Combat correction as suggested by Kolodziejczyk et al^38^.

### Multidimensional scaling analysis (MDS)

A number of factors could be invoked to explain the two groups observed in our multidimensional scaling analysis (MDS) (Figure 1B), but they should also explain that only one group is structured according to developmental time. The obvious hypothesis that they correspond to male and female embryos does not explain that structure, and is also not supported by examining X/autosome gene expression ratios (Figure S16). An alternative hypothesis is that samples in the small cluster are unfertilized eggs. If an egg is not fertilized, after completion of meiosis, development will be arrested^39^, but they are visually indistinguishable. This hypothesis is confirmed by at least two lines of evidence, in addition to the lack of developmental time structure. First, for expression correlation, all samples in the small cluster are highly correlated with unfertilized egg, while the correlations in the other samples gradually decrease with development (Figure S3A). Second, all the samples from the small cluster are enriched with meiosis related genes (Figure S3B). Thus we excluded the small cluster for downstream analyses, i.e. we used 150 embryos with an average of 18 individuals per developmental stage.

### Metrics of expression variability

Expression variability is generally measured by the coefficient of variation (CV)^40^. However, a gene’s CV is strongly dependent on its RNA abundance (Figure S4). While this is an inherent property of time-interval counting processes (such as a Poisson process), it makes the comparison of variability between different conditions difficult^38,41^. Distance to median (DM, the distance between the squared CV of a gene and its running median) has been proposed as a variability metric that is independent of expression level^38,41,42^. However, the DM is still strongly negatively correlated with the mean expression level in our data (Figure S5). To avoid this dependency, we developed another variability measure, the adjusted standard deviation (adjusted SD), by calculating the ratio between observed SD and expected SD. Following the same approach as Barroso et al.^43^, we performed polynomial regressions to predict the expected SD from mean expression. Since the adjusted SD metric works much better than DM in terms of accounting for the confounding effects of mean expression (Figure S6), we adopted it as a measure of expression variability in our study. As observed in yeast^42,44^, we found that essential genes and hubs (proteins in the center of protein-protein interaction network) have lower expression variability than other genes (Figure S17), indicating selection to reduce it. This observation provides a control that we are indeed measuring biologically relevant expression variability.

Detailed calculation of expression variability:

#### 1. Adjusted SD

For each gene, we computed standard deviation (SD) in each stage and over all stages. Then we fitted a polynomial model to predict the global (across all stages) SD from the global mean expression. We increased the degrees of the model until there was no more significant improvement (tested with ANOVA, *p*<0.05 as a significant improvement). Then, based on this best fitting model, for each gene, we computed its predicted global SD based on its global mean expression. Finally, the adjusted SD of a gene in one stage is this gene’s SD in its corresponding stage divided by its predicted global SD. This method is derived from Barroso et al.^43^, but computing adjusted SD rather than adjusted variance.

#### 2. Distance to median: the distance between the squared coefficient of variation (CV) of a gene and its running median

For each gene, we computed its squared CV in each stage and over all stages. Then, we ordered genes based on their global (across all stages) mean expression. Next, we defined series of sliding windows of 50 genes with 25 genes overlap, starting from lowest global mean expression. Finally, the distance to median of a gene in one stage is the stage specific log10 squared CV minus the median of global log10 squared CV in this gene’s corresponding window. R code was modified from the DM function of the scran package^34^.

### Bootstrap analysis

For each stage, we randomly sampled the same number of samples. Then, we computed the adjusted SD based on these random samples. We repeated the first two steps 500 times. Each time, we only kept the median of the adjusted SD for each stage. Thus in each stage we obtained 500 medians. Finally, we performed a Wilcoxon test to test the significance of the difference between the bootstrapped values of different stages.

### ChIP-Seq data analysis

#### 1. Histone modification signal datasets

The signal data files of four euchromatin histone modification marks (H3K4me1, H3K4me3, H3K9ac, and H3K27ac) at six developmental stages (0-4h, 4-8h, 8-12h, 12-16h. 16-20h, 20-24h) were downloaded from modENCODE^26^ (NCBI GEO: GSE16013) (March, 2018). The signal is smoothed, background-subtracted tag density. The signal was precomputed along the genome in 35-bp windows.

#### 2. Histone modification signal for promoter and gene body

For each gene, as suggested by Nicolas et al.^23^, we separately calculated the mean signal of its proximal promoter (2 kb upstream to 2 kb downstream for transcription start site (TSS)) and of its gene body (TSS to transcription end site (TES)) by using the bedtools “map” command^45^. The TSS and TES information was retrieved from Ensembl release 91^32^. For a gene with several TSS and TES, we use its mean coordinates.

#### 3. Histone modification signal Z score transformation

For each modification mark in each stage, the signal value was transformed into a Z score by subtracting the mean signal across intergenic regions and dividing by the standard deviation signal of the intergenic regions. The intergenic region were defined by removing all proximal promoter regions and gene body regions with the bedtools “subtract” command^45^. Our assumption is that on average such intergenic regions are not the target of active histone modification signal, and thus allow to normalize between libraries. Then, for each gene, we re-calculated the mean signal (Z score) of its proximal promoter (2 kb upstream to 2 kb downstream for transcription start site (TSS)) and of its gene body (TSS to transcription end site (TES)) by using the bedtools “map” command^45^.

### Identification of stage specifically expressed genes

Following the same approach as previously^46^, we first defined 8 stage specific expressed artificial expression profile (Figure S18A). Then, for each gene, we performed Pearson’s correlation between its real expression and this artificial expression. Finally, for each artificial expression profile, we kept genes with top 10% correlation coefficient as the corresponding stage specifically expressed genes (Figure S18B).

### Identification of hourglass expression variability genes

Similar to the stage specifically expressed gene identification approach, we correlated each gene’s variability profile with the median across all genes. Then, we kept genes with the top 10% correlation coefficient as the ones following the global hourglass variability profile.

### Identification of genes expressed at all stages

For each gene, we calculated the average expression across replicates in each stage. Then, we defined the average expression > 1 as expressed.

### Identification of genes with constant expression across all stages

For each gene, we first preformed one-way analysis of variance (ANOVA) to compare the means of expression in different stage. Then, we calculated the *q-values* for multiple test correction. Finally, the constantly expressed genes were defined as genes with *q-values* > 0.05.

### Gene ontology (GO) enrichment analysis

We performed GO enrichment analysis for hourglass expression variability genes by using the topGO^47^ R package with the “elim” algorithm.

### Single Nucleotide Polymorphism (SNP) data

The SNP data for 205 *D. melanogaster* inbred lines were downloaded from the Drosophila Genetic Reference Panel (DGRP^48^) (December, 2018).

### Nucleotide diversity (π) calculation

We calculated nucleotide diversity of promoters with vcftools^49^.

### Transcriptome index analysis

A “transcriptome index”^50,51^ is a weighted mean of a feature over all genes, where the weights are the expression levels of the genes at each condition (e.g., developmental stage). The transcriptome index of phastCons was calculated as:

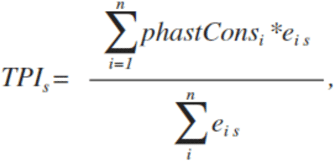

where *s* is the developmental stage, *phastCons*_*i*_ is the promoter sequence conservation score of gene *i*, *n* is the total number of genes, and *e*_*is*_ is the expression level (log transformed) of gene *i* in developmental stage *s*. For the transcriptome index of nucleotide diversity (π) the same formula is used, replacing *phastCons*_*i*_ by *π*_*i*_.

### Meiosis related genes and transcription factors

The Meiosis related genes and transcription factors were downloaded from AmiGO^52^ (May, 2018).

### Individual unfertilized eggs RNA-seq data

The normalized and log transformed expression matrix of individual unfertilized eggs was downloaded from NCBI GEO: GSE68062^53^ (May, 2018).

### Dispersed and precise promoters

The annotation of genes with dispersed or precise promoters was downloaded from Schor et al^20^ (June, 2019). Dispersed promoters are often associated with ubiquitously expressed genes, have more dispersed patterns of transcriptional initiation, and do not contain a TATA box. On the contrary, precise promoters are typically associated with restricted tissue-specific expression and with a TATA box, and have a single predominant TSS.

### Essential gene annotation and protein connectivity datasets

The gene essentiality and protein connectivity datasets were downloaded from OGEE v2^54^ (March, 2018).

### PhastCons score

The pre-computed sequence conservation score phastCons^55^ of fly genome was downloaded from http://hgdownload.soe.ucsc.edu/goldenPath/dm3/phastCons15way/ (February, 2018). Higher value means higher conservation.

### Experimentally validated core promoters

Experimentally validated transcription start sites (TSSs) were downloaded from the Eukaryotic Promoter Database (EPD)^29^ (May, 2018). For a gene with several TSSs, we selected the most representative one (the TSS that has been validated by the largest number of samples). The core promoter region was defined as 49 bp upstream TSS to 10 bp downstream of the TSS^29^. We used EPD defined TSSs here because they are more accurate for defining core promoters, whose function is expected to be related to sequence conservation. Whereas for histone modification signal for promoter and gene body we used Ensembl defined TSSs to be consistent with the source of TES information, and precision was less important in defining broader proximal promoters.

## Acknowledgements

We thank Virginie Braman for help with embryo collection and library preparation. We thank Daniel Alpern for help with the BRB-seq technology. We thank Richard Benton, David Garfield and Laurent Keller for critically reading and commenting on the manuscript. We thank Andrea Komljenovic and other members of the Robinson-Rechavi lab for helpful discussions. Part of the computations were performed at the Vital-IT (http://www.vital-it.ch) Center for high-performance computing of the SIB Swiss Institute of Bioinformatics. JL and MRR are supported by Swiss National Science Foundation grants 31003A_153341 / 1 and 31003A_173048. MRR and BD are supported by SystemsX.ch grant AgingX.

## Author contributions

JL designed the work with input from MRR, MF and BD. MF led all experiments. JL performed all computational analyses. VG and BD contributed expertise in the BRB-seq experiments. JL and MRR interpreted the results with input from all the other authors. JL wrote the first draft of the paper. JL and MRR finalized the paper with input from all the other authors.

## Supplementary figure legends

**Figure S1: Relationship between uniquely mapped reads and expressed genes**

Each dot represents one sample. The black dots indicate low quality samples with <4500 expressed genes or with <0.3 million uniquely mapped reads. The 239 orange colored samples were retained for downstream analysis (“high quality samples”).

**Figure S2: Proportion of retained samples in each development stage**

The number of retained samples and of total samples in each stage is indicated in the bottom of each bar.

**Figure S3: Evidence that the samples from the small cluster are unfertilized eggs**

A. Boxplot of Spearman’s correlation coefficients (rho) of expression between individual unfertilized eggs and each sample from the small cluster or from the large cluster, showing that the small cluster has an expression profile of unfertilized eggs. The lower and upper intervals indicated by the dashed lines (“whiskers”) represent 1.5 times the interquartile range (IQR), and the box shows the lower and upper intervals of IQR together with the median.

B. Expression heat map of meiosis related genes across all samples, showing that their expression decreases over development for the large cluster, but is high in all samples of the small cluster, consistent with unfertilized eggs.

For testing of an alternative explanation of the two clusters as being males and females, see Figure S16.

**Figure S4: Relationship between average expression and coefficient of variation at each stage**

Pearson’s correlation between average expression and coefficient of variation in each development stage is indicated in the top left of each subfigure.

**Figure S5: Relationship between average expression and distance to median at each stage**

Pearson’s correlation between average expression and distance to median in each development stage is indicated in the top left of each subfigure.

**Figure S6: Relationship between average expression and adjusted SD at each stage**

Pearson’s correlation between average expression and adjusted SD in each development stage is indicated in the top left of each subfigure.

**Figure S7: Variation of expression variability across development using alternate measures of variability**

A. Variability measured by adjusted SD; unlike in Figure 2, the variability in E1 was calculated using all samples from both small and large clusters.

B. Variability measured by coefficient of variation (CV).

C. Variability measured by distance to median (DM).

The legend is the same as for Figure 2. We performed pairwise Wilcoxon test between any two stages to test the significance. The multiple test corrected *p*-values (Benjamini–Hochberg method) are shown in Tables S6, S7 and S8.

**Figure S8: Bootstrap analysis of the variability calculation** We performed pairwise Wilcoxon test between any two stages to test the significance. The multiple test corrected *p*-values (Benjamini–Hochberg method) are shown in Table S9.

**Figure S9: Variation of expression variability across development for different categories of genes**

A. Genes with constant expression level over development.

B. Transcription factor.

The legend is the same as for Figure 2. We performed pairwise Wilcoxon test between any two stages to test the significance. The multiple test corrected *p*-values (Benjamini–Hochberg method) are shown in Tables S10 and S11.

**Figure S10: Variation of expression variability across development for dispersed promoter genes and for precise promoter genes**

For each stage, the first and the second box represents dispersed promoter genes and precise promoter genes respectively. The legend is the same as for Figure 2. We performed pairwise Wilcoxon test between any two stages to test the significance separately for dispersed promoter genes and for precise promoter genes. The multiple test corrected *p*-values (Benjamini– Hochberg method) are shown in Tables S12 and S13.

**Figure S11: Histone modification signal across development**

The legend is the same as for Figure 3B and 3C. The median signal value in each development stage is indicated above each box. We performed pairwise Wilcoxon test between any two stages to test the significance. The multiple test corrected *p*-values (Benjamini–Hochberg method) for H3K4Me1, H3K27Ac and H3K9Ac are shown in Tables S14-S19.

**Figure S12: Spearman’s correlation coefficient between histone modification signal and promoter nucleotide diversity (π).**

The legend is the same as for Figure 4A.

**Figure S13: Spearman’s correlation coefficient between histone modification signal and promoter sequence conservation for different definitions of promoter width**

The figure legend is the same as in Figure 4A.

A. Promoter defined as 200 bp around TSS

B. Promoter defined as 400 bp around TSS

C. Promoter defined as 1000 bp around TSS

**Figure S14: transcriptome index of π across development.**

The legend is the same as for Figure 4C.

**Figure S15: Multidimensional scaling analysis for all samples**

Different colors indicate different stages. The solid triangles represent high quality samples according to Figure S1; the hollow triangles represent low quality samples which were discarded.

**Figure S16: Mapping of X/autosome gene expression ratios to the multidimensional scaling analysis plot**

We calculated the ratio of mean expression between genes from the X chromosome and from the autosomes for each sample. Red represents high ratio, blue represents low ratio. For Drosophila, dosage compensation is achieved by increasing expression of X chromosome genes in males. Since the dosage compensation is still incomplete during development, females should have a higher ratio of mean expression between genes from the X chromosome and from the autosomes. Here, we found both high ratio samples and low ratio samples are well mixed in both the cluster and large clusters. Thus, we reject the hypothesis that the two different clusters are due to sex.

**Figure S17: Relationship between expression variability and protein importance**

We used the average variability across all development stages.

A. We split genes into 10 equally sized bins based on expression variability. The proportion of essential genes was fit by regression (the first degree of polynomial), whose *R*^2^ and *p*-value are indicated in the top-left corner of each graph. The median expression variability of each bin was plotted on the x-axis.

B. Spearman’s correlation between connectivity in a protein-protein interaction network and expression variability. The coefficient and *p*-value are indicated in the top-right. Loess regression lines are plotted in red.

**Figure S18: Detection of stage specific genes**

A. The artificial expression profile.

B. The expression of identified stage specific genes. The bold black line represents the median expression, the two gray lines represent 25th and 75th quantiles of expression, respectively.

