## Supplementary Figures S1-S18 for "Selection against expression noise explains the origin of the hourglass pattern of Evo-Devo"

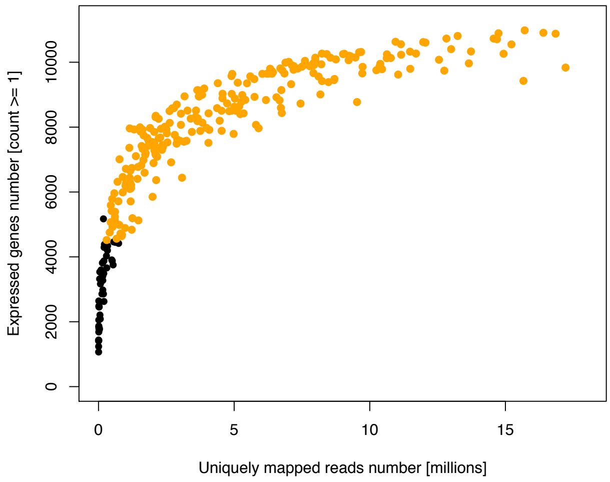


***Figure S1: Relationship between uniquely mapped reads and expressed genes***

*Each dot represents one sample. The black dots indicate low quality samples with <4500 expressed genes or with <0.3 million uniquely mapped reads. The 239 orange colored samples were retained for downstream analysis ("high quality samples").*

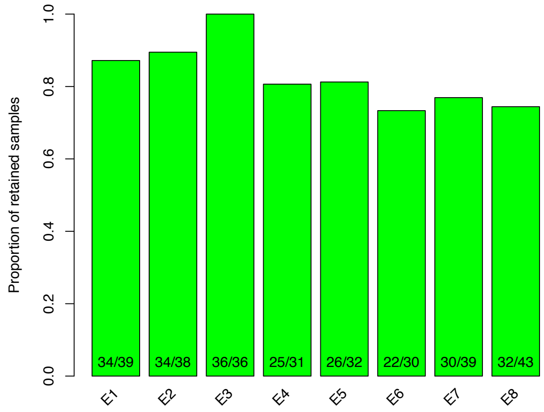


***Figure S2: Proportion of retained samples in each development stage***

*The number of retained samples and of total samples in each stage is indicated in the bottom of each bar.*

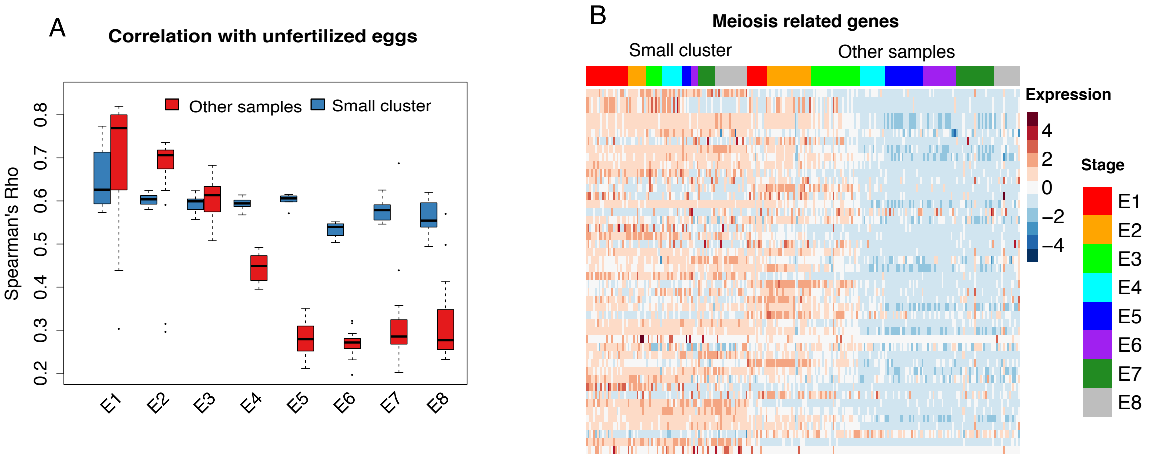


***Figure S3: Evidence that the samples from the small cluster are unfertilized eggs***

A. *Boxplot of Spearman's correlation coefficients (rho) of expression between individual unfertilized eggs and each sample from the small cluster or from the large cluster, showing that the small cluster has an expression profile of unfertilized eggs. The lower and upper intervals indicated by the dashed lines ("whiskers") represent 1.5 times the interquartile range (IQR), and the box shows the lower and upper intervals of IQR together with the median.*

B. *Expression heat map of meiosis related genes across all samples, showing that their expression decreases over development for the large cluster, but is high in all samples of the small cluster, consistent with unfertilized eggs.*

*For testing of an alternative explanation of the two clusters as being males and females, see Figure S16.*

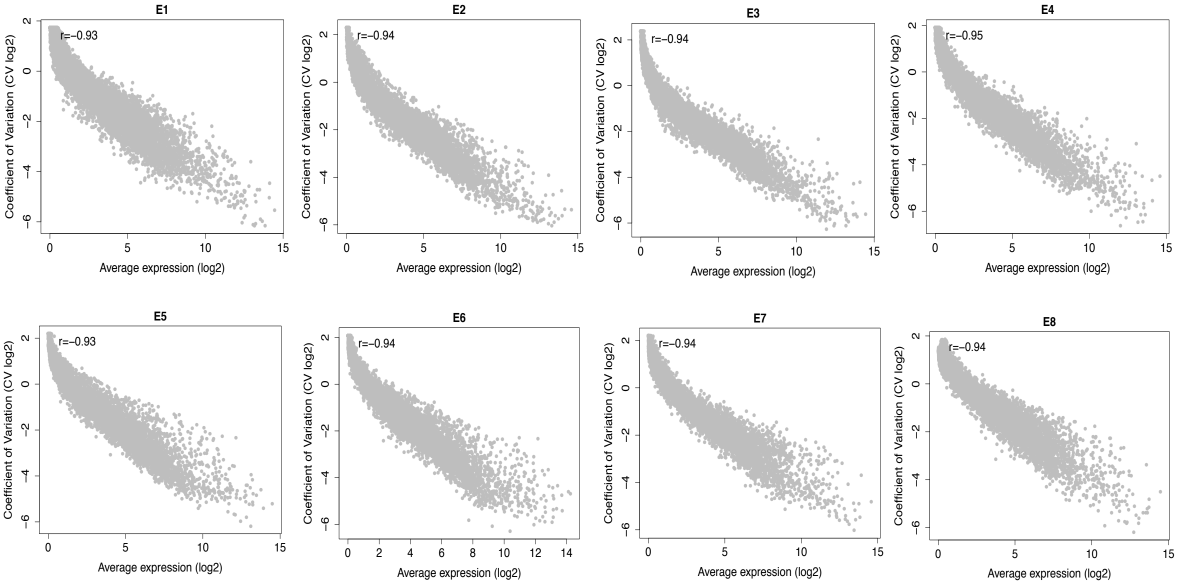


***Figure S4: Relationship between average expression and coefficient of variation at each stage***

*Pearson's correlation between average expression and coefficient of variation in each development stage is indicated in the top left of each subfigure.*

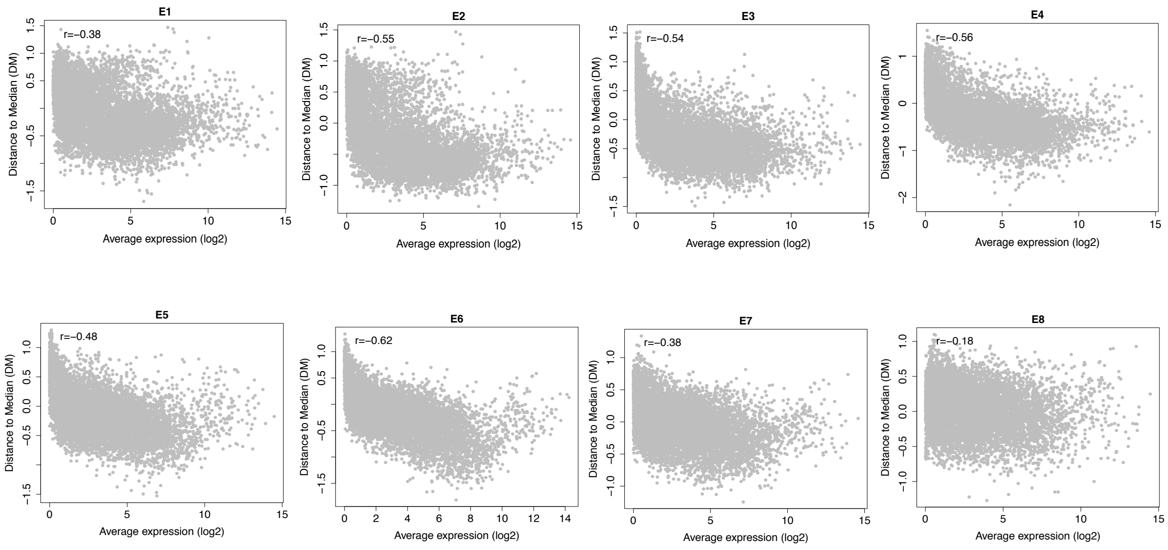


***Figure S5: Relationship between average expression and distance to median at each stage***

*Pearson's correlation between average expression and distance to median in each development stage is indicated in the top left of each subfigure.*

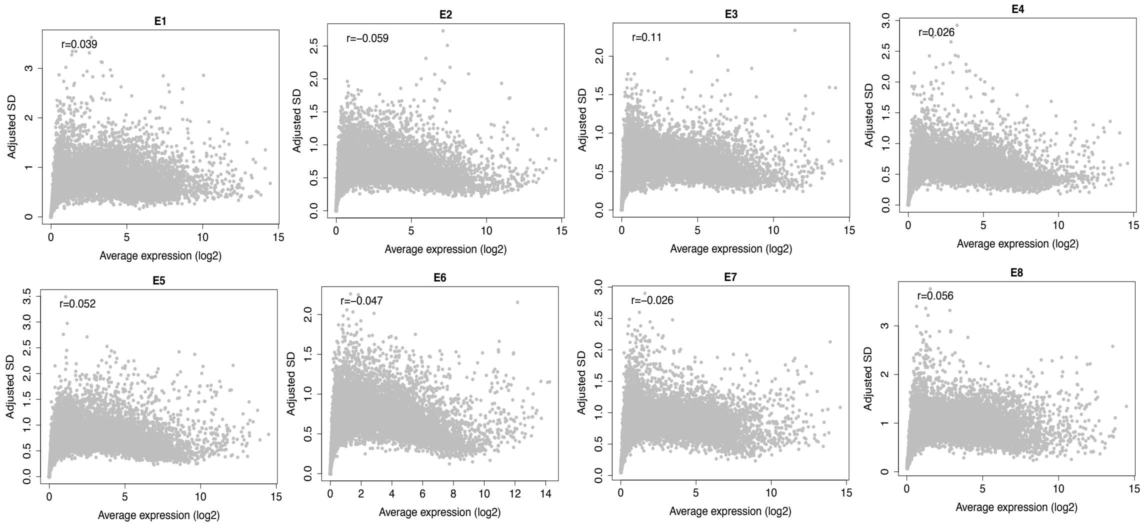


***Figure S6: Relationship between average expression and adjusted SD at each stage***

*Pearson's correlation between average expression and adjusted SD in each development stage is indicated in the top left of each subfigure.*

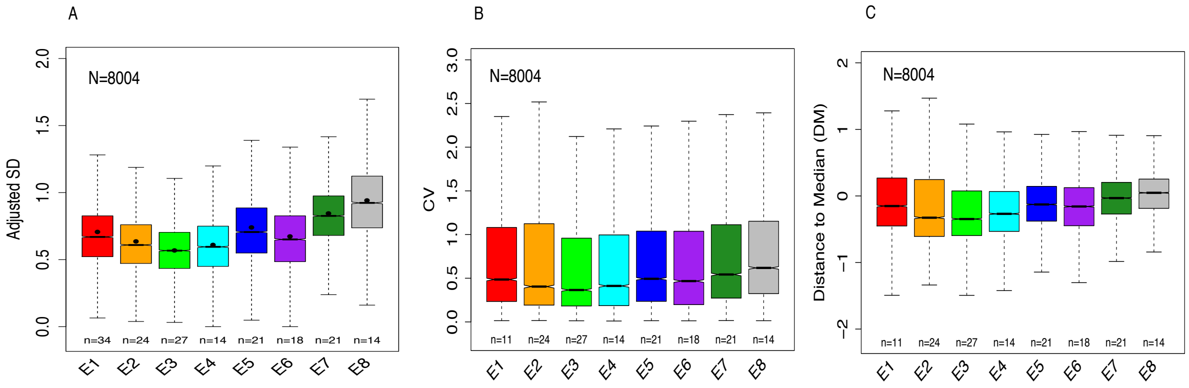


***Figure S7: Variation of expression variability across development using alternate measures of variability***

*A. Variability measured by adjusted SD; unlike in Figure 2, the variability in E1 was calculated using all samples from both small and large clusters.*

*B. Variability measured by coefficient of variation (CV).*

*C. Variability measured by distance to median (DM).*

*The legend is the same as for Figure 2. We performed pairwise Wilcoxon test between any two stages to test the significance. The multiple test corrected p-values (Benjamini–Hochberg method) are shown in Tables S6, S7 and S8.*

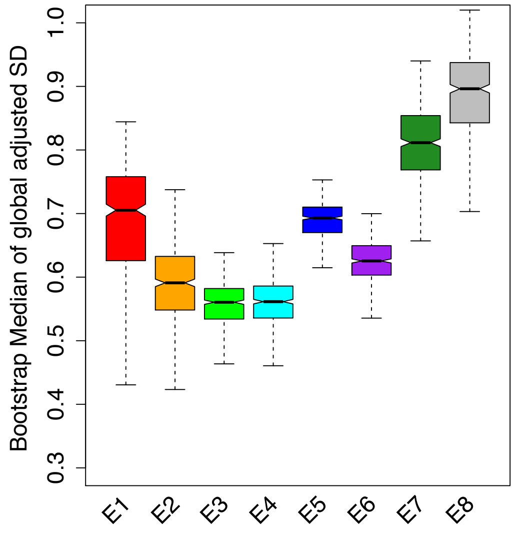


***Figure S8: Bootstrap analysis of the variability calculation***

*We performed pairwise Wilcoxon test between any two stages to test the significance. The multiple test corrected p-values (Benjamini–Hochberg method) are shown in Table S9.*

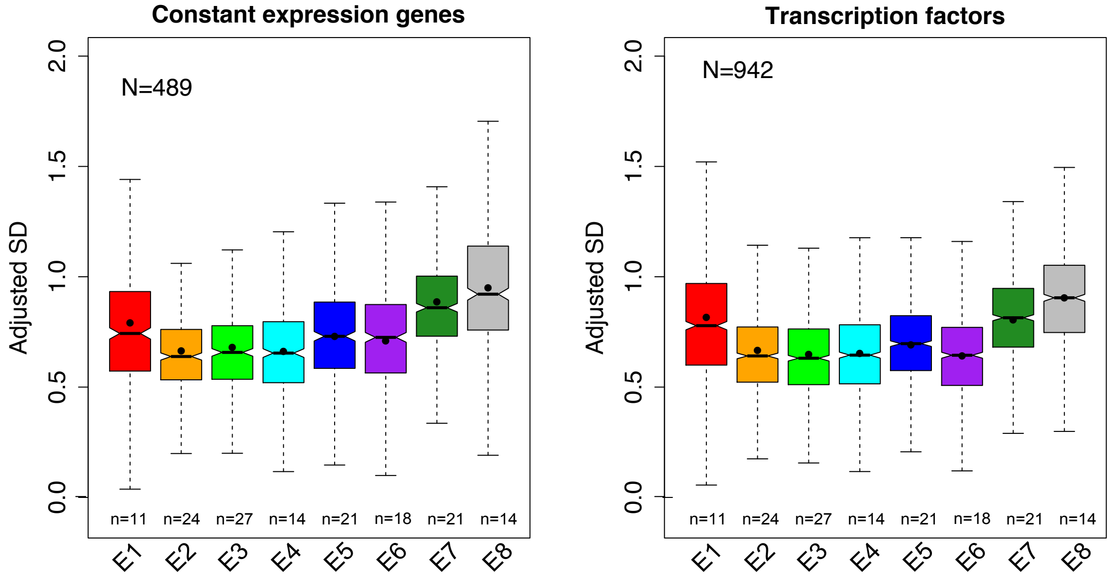


***Figure S9: Variation of expression variability across development for different categories of genes***

*A. Genes with constant expression level over development.*

*B. Transcription factor.*

*The legend is the same as for Figure 2. We performed pairwise Wilcoxon test between any two stages to test the significance. The multiple test corrected p-values (Benjamini–Hochberg method) are shown in Tables S10 and S11.*

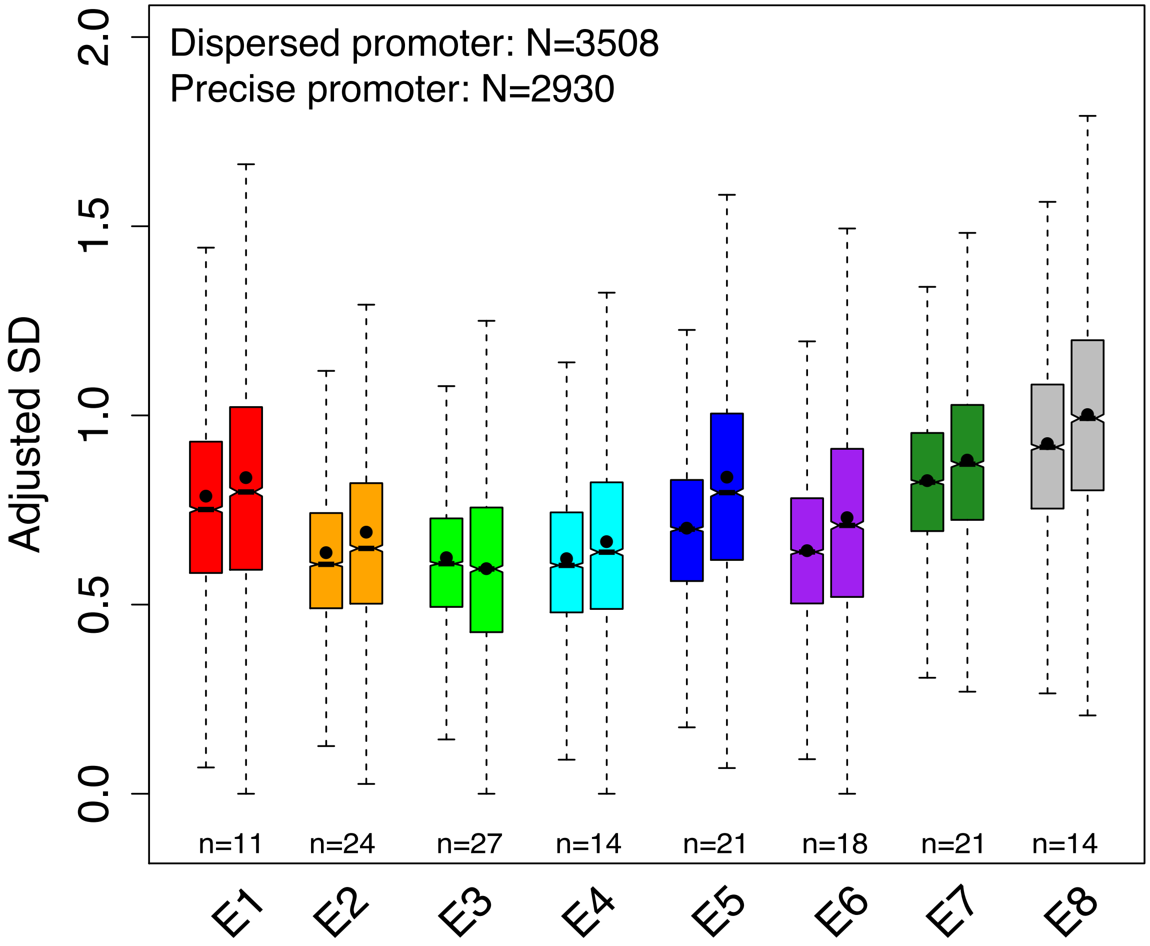


***Figure S10: Variation of expression variability across development for dispersed promoter genes and for precise promoter genes***

*For each stage, the first and the second box represents dispersed promoter genes and precise promoter genes respectively. The legend is the same as for Figure 2. We performed pairwise Wilcoxon test between any two stages to test the significance separately for dispersed promoter genes and for precise promoter genes. The multiple test corrected p-values (Benjamini–Hochberg method) are shown in Tables S12 and S13.*

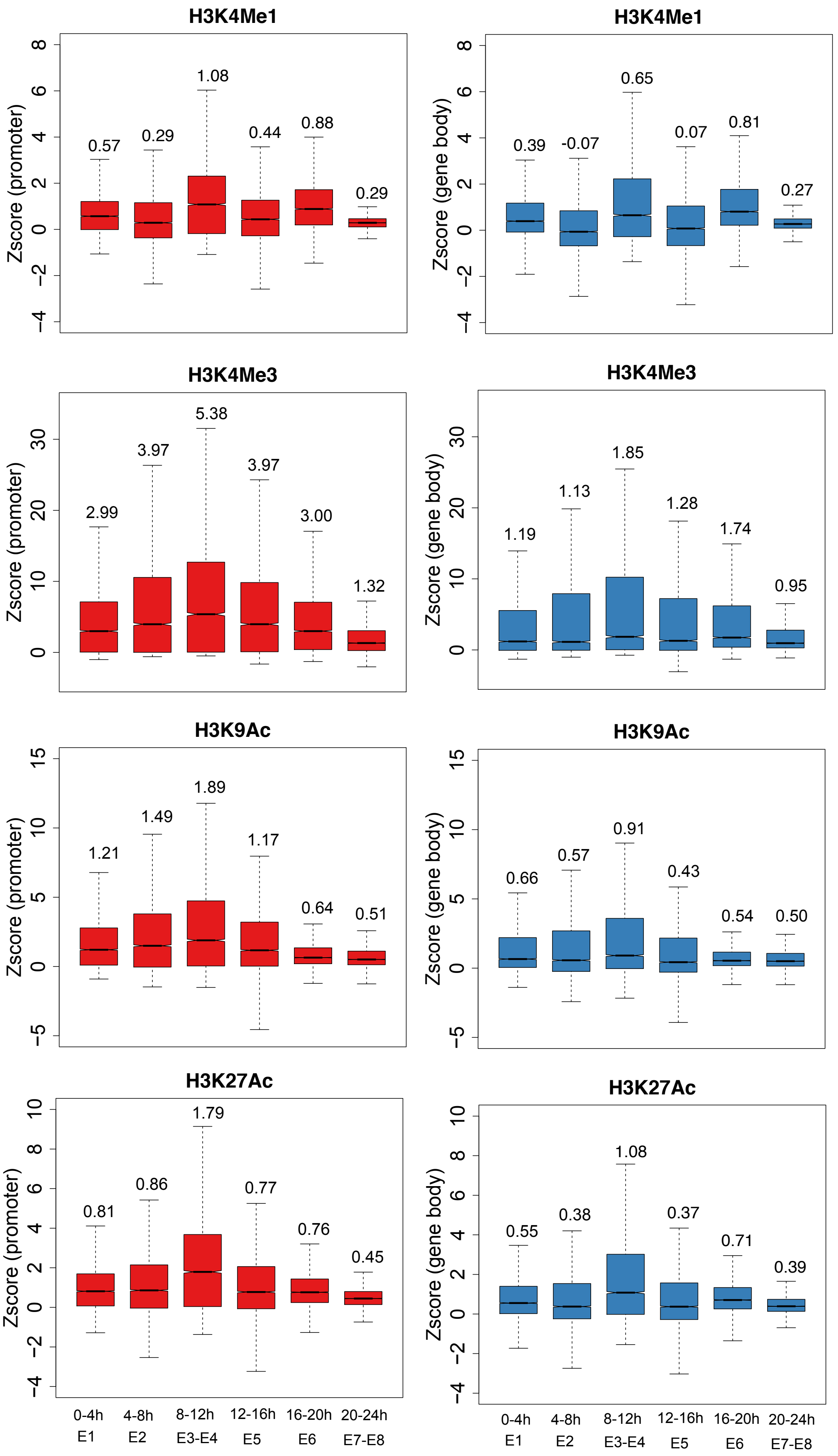
H3K4Me1
Zscore (promoter)
0.57
0.29
1.08
0.44
0.88
0.29
H3K4Me1
Zscore (gene body)
0.39
-0.07
0.65
0.07
0.81
0.27
H3K4Me3
Zscore (promoter)
2.99
3.97
5.38
3.97
3.00
1.32
H3K4Me3
Zscore (gene body)
1.19
1.13
1.85
1.28
1.74
0.95
H3K9Ac
Zscore (promoter)
1.21
1.49
1.89
1.17
0.64
0.51
H3K9Ac
Zscore (gene body)
0.66
0.57
0.91
0.43
0.54
0.50
H3K27Ac
Zscore (promoter)
0.81
0.86
1.79
0.77
0.76
0.45
H3K27Ac
Zscore (gene body)
0.55
0.38
1.08
0.37
0.71
0.39
0-4h
E1
4-8h
E2
8-12h
E3-E4
12-16h
E5
16-20h
E6
20-24h
E7-E8

***Figure S11: Histone modification signal across development***

*The legend is the same as for Figure 3B and 3C. The median signal value in each development stage is indicated above each box. We performed pairwise Wilcoxon test between any two stages to test the significance. The multiple test corrected p-values (Benjamini–Hochberg method) for H3K4Me1, H3K27Ac and H3K9Ac are shown in Tables S14-S19.*

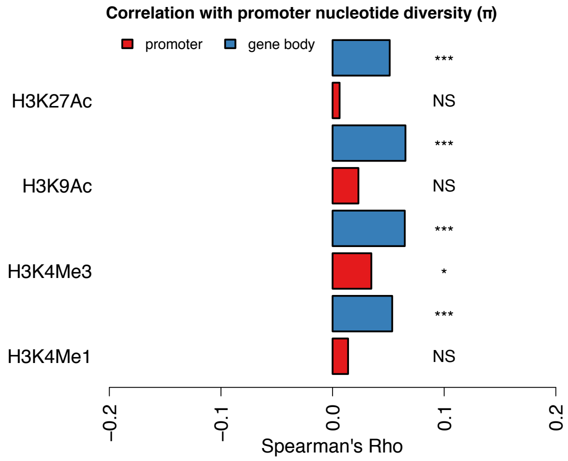


***Figure S12: Spearman's correlation coefficient between histone modification signal and promoter nucleotide diversity (π).***

*The legend is the same as for Figure 4A.*

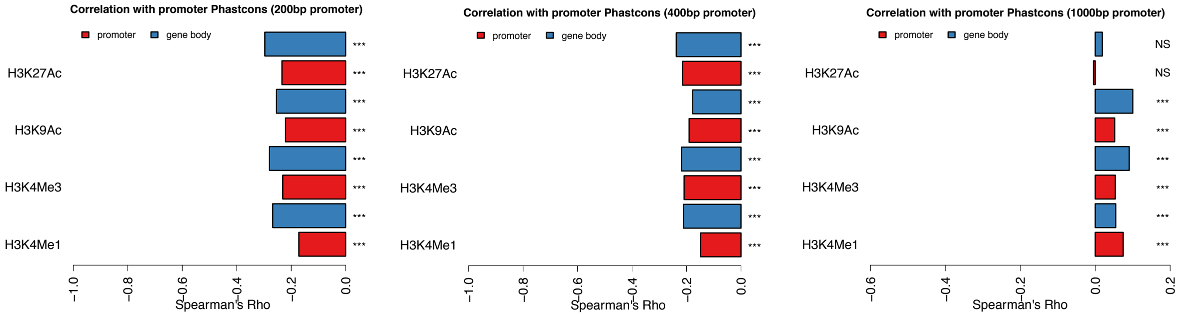


***Figure S13: Spearman's correlation coefficient between histone modification signal and promoter sequence conservation for different definitions of promoter width***

*The figure legend is the same as in Figure 4A.*

*A. Promoter defined as 200 bp around TSS*

*B. Promoter defined as 400 bp around TSS*

*C. Promoter defined as 1000 bp around TSS*

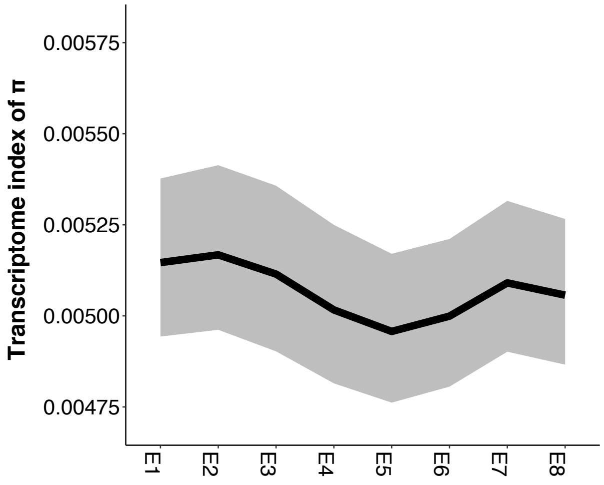


***Figure S14: transcriptome index of π across development.***

*The legend is the same as for Figure 4C.*

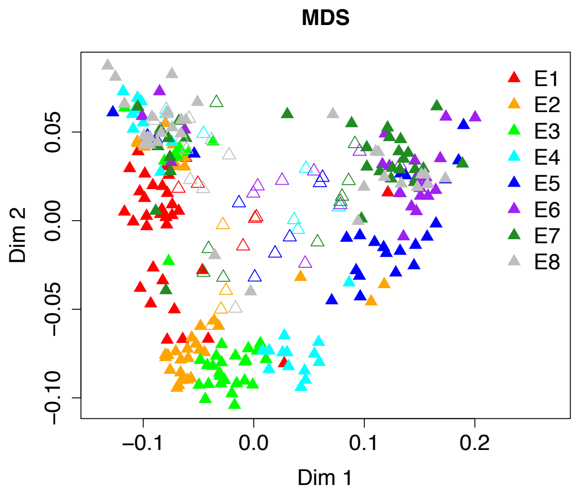


***Figure S15: Multidimensional scaling analysis for all samples***

*Different colors indicate different stages. The solid triangles represent high quality samples according to Figure S1; the hollow triangles represent low quality samples which were discarded.*

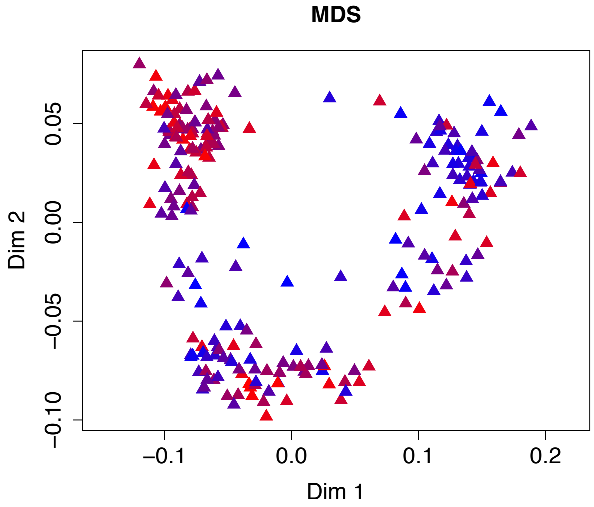


***Figure S16: Mapping of X/autosome gene expression ratios to the multidimensional scaling analysis plot***

*We calculated the ratio of mean expression between genes from the X chromosome and from the autosomes for each sample. Red represents high ratio, blue represents low ratio. For Drosophila, dosage compensation is achieved by increasing expression of X chromosome genes in males. Since the dosage compensation is still incomplete during development, females should have a higher ratio of mean expression between genes from the X chromosome and from the autosomes. Here, we found both high ratio samples and low ratio samples are well mixed in both the cluster and large clusters. Thus, we reject the hypothesis that the two different clusters are due to sex.*

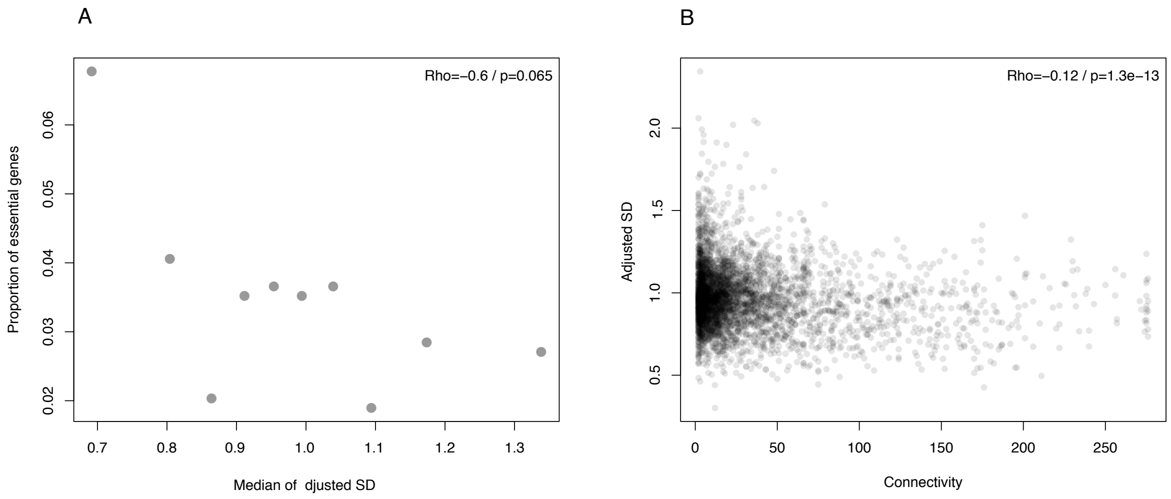


***Figure S17: Relationship between expression variability and protein importance***

*We used the average variability across all development stages.*

A. *Spearman's correlation between expression variability and proportion of essential genes. We split genes into 10 equally sized bins based on expression variability. The Spearman's correlation is calculated by using the median variability and proportion of essential genes in the ten bins. The Spearman's correlation coefficient and p-value are indicated in the top-right.*

B. *Spearman's correlation between connectivity in a protein-protein interaction network and expression variability. The coefficient and p-value are indicated in the top-right.*

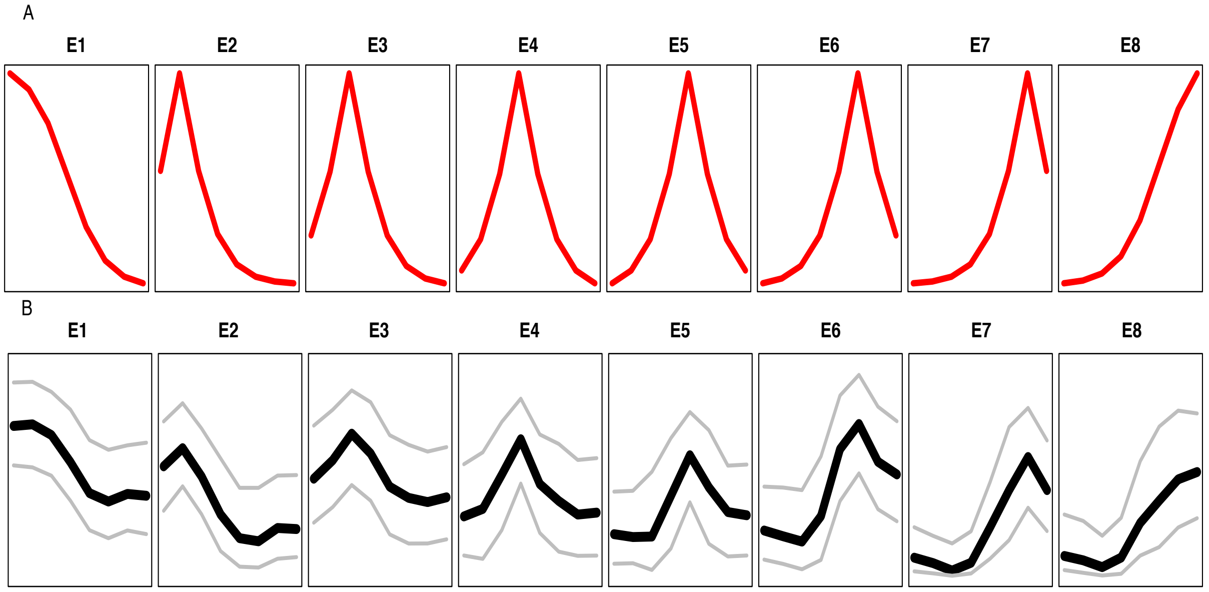


***Figure S18: Detection of stage specific genes***

A. *The artificial expression profile.*

*B. The expression of identified stage specific genes. The bold black line represents the median expression, the two gray lines represent 25th and 75th quantiles of expression, respectively.*
