## Supplementary Tables S1-S19 for "Selection against expression noise explains the origin of the hourglass pattern of Evo-Devo"

**Table S1: variability (adjusted SD) comparison between any two stages (all genes)**

Wilcoxon test, multiple test corrected *p*-values (Benjamini–Hochberg method).

| Stages | E1 | E2 | E3 | E4 | E5 | E6 | E7 |
| --- | --- | --- | --- | --- | --- | --- | --- |
| E2 | 3.11e-180 |  |  |  |  |  |  |
| E3 | 0.00e+00 | 6.17e-43 |  |  |  |  |  |
| E4 | 1.21e-247 | 5.24e-08 | 2.49e-16 |  |  |  |  |
| E5 | 1.06e-13 | 2.57e-118 | 4.14e-292 | 1.99e-175 |  |  |  |
| E6 | 1.93e-92 | 3.99e-17 | 2.62e-105 | 8.75e-43 | 1.14e-44 |  |  |
| E7 | 2.57e-78 | 0.00e+00 | 0.00e+00 | 0.00e+00 | 3.06e-181 | 0.00e+00 |  |
| E8 | 6.64e-268 | 0.00e+00 | 0.00e+00 | 0.00e+00 | 0.00e+00 | 0.00e+00 | 3.14e-106 |

**Table S2: variability (adjusted SD) comparison between any two stages (genes expressed at all stages)**

Wilcoxon test, multiple test corrected *p*-values (Benjamini–Hochberg method).

| Stages | E1 | E2 | E3 | E4 | E5 | E6 | E7 |
| --- | --- | --- | --- | --- | --- | --- | --- |
| E2 | 8.57e-175 |  |  |  |  |  |  |
| E3 | 3.04e-254 | 2.92e-06 |  |  |  |  |  |
| E4 | 6.57e-170 | 2.39e-01 | 7.21e-10 |  |  |  |  |
| E5 | 2.05e-17 | 3.71e-125 | 1.92e-200 | 7.95e-113 |  |  |  |
| E6 | 9.89e-105 | 1.45e-17 | 2.93e-45 | 2.38e-13 | 2.87e-52 |  |  |
| E7 | 9.01e-47 | 0.00e+00 | 0.00e+00 | 0.00e+00 | 1.97e-163 | 0.00e+00 |  |
| E8 | 3.11e-199 | 0.00e+00 | 0.00e+00 | 0.00e+00 | 0.00e+00 | 0.00e+00 | 2.19e-107 |

**Table S3: proximal promoter H3K4Me3 signal (Z score) comparison between any two stages**

Wilcoxon test, multiple test corrected *p*-values (Benjamini–Hochberg method).

| Satges | 0-4h | 4-8h | 8-12h | 12-16h | 16-20h |
| --- | --- | --- | --- | --- | --- |
| 4-8h | 2.39e-36 |  |  |  |  |
| 8-12h | 3.71e-93 | 5.51e-13 |  |  |  |
| 12-16h | 6.35e-35 | 6.84e-01 | 1.28e-17 |  |  |
| 16-20h | 2.29e-09 | 6.05e-08 | 4.17e-47 | 7.73e-11 |  |
| 20-24h | 3.10e-88 | 2.08e-118 | 8.88e-197 | 1.40e-161 | 9.07e-182 |

**Table S4: gene body H3K4Me3 signal (Z score) comparison between any two stages**

| Satges | 0-4h | 4-8h | 8-12h | 12-16h | 16-20h |
| --- | --- | --- | --- | --- | --- |
| 4-8h | 1.72e-12 |  |  |  |  |
| 8-12h | 2.96e-63 | 2.45e-23 |  |  |  |
| 12-16h | 6.29e-06 | 2.89e-02 | 3.57e-31 |  |  |
| 16-20h | 2.56e-51 | 2.46e-21 | 2.55e-01 | 2.45e-22 |  |
| 20-24h | 2.55e-01 | 9.06e-04 | 1.08e-45 | 2.79e-06 | 1.03e-89 |

**Table S5: promoter sequence conservation comparison between any two stages**

Wilcoxon test, multiple test corrected *p*-values (Benjamini–Hochberg method).

| Stages | E1 | E2 | E3 | E4 | E5 | E6 | E7 |
| --- | --- | --- | --- | --- | --- | --- | --- |
| E2 | 5.02e-01 |  |  |  |  |  |  |
| E3 | 1.70e-11 | 4.66e-12 |  |  |  |  |  |
| E4 | 5.57e-29 | 1.96e-28 | 4.06e-06 |  |  |  |  |
| E5 | 1.07e-35 | 7.75e-35 | 2.93e-10 | 8.85e-02 |  |  |  |
| E6 | 2.18e-37 | 3.23e-36 | 5.29e-09 | 4.19e-01 | 3.13e-01 |  |  |
| E7 | 1.89e-11 | 5.11e-12 | 7.91e-01 | 2.09e-05 | 2.13e-09 | 8.87e-08 |  |
| E8 | 2.28e-21 | 1.72e-21 | 2.18e-02 | 9.67e-03 | 7.81e-06 | 1.79e-04 | 6.34e-02 |

**Table S6: variability (adjusted SD) comparison between any two stages**

Here, the E1 stage contains all samples from both large and small clusters. Wilcoxon test, multiple test corrected *p*-values (Benjamini–Hochberg method).

| Stages | E1 | E2 | E3 | E4 | E5 | E6 | E7 |
| --- | --- | --- | --- | --- | --- | --- | --- |
| E2 | 5.49e-58 |  |  |  |  |  |  |
| E3 | 6.24e-202 | 2.55e-46 |  |  |  |  |  |
| E4 | 1.02e-96 | 1.57e-07 | 1.60e-18 |  |  |  |  |
| E5 | 9.25e-19 | 1.32e-124 | 3.41e-303 | 6.09e-175 |  |  |  |
| E6 | 2.56e-09 | 3.21e-19 | 6.02e-112 | 2.66e-43 | 1.86e-43 |  |  |
| E7 | 0.00e+00 | 0.00e+00 | 0.00e+00 | 0.00e+00 | 1.56e-163 | 0.00e+00 |  |
| E8 | 0.00e+00 | 0.00e+00 | 0.00e+00 | 0.00e+00 | 0.00e+00 | 0.00e+00 | 6.72e-104 |

**Table S7: variability (coefficient of variation) comparison between any two stages**

Wilcoxon test, multiple test corrected *p*-values (Benjamini–Hochberg method).

| Stages | E1 | E2 | E3 | E4 | E5 | E6 | E7 |
| --- | --- | --- | --- | --- | --- | --- | --- |
| E2 | 9.30e-9 |  |  |  |  |  |  |
| E3 | 1.35e-25 | 5.29e-05 |  |  |  |  |  |
| E4 | 1.24e-15 | 3.32e-02 | 4.33e-02 |  |  |  |  |
| E5 | 6.79e-01 | 3.64e-08 | 1.22e-24 | 9.45e-15 |  |  |  |
| E6 | 1.70e-05 | 1.94e-01 | 1.69e-08 | 3.31e-04 | 9.93e-05 |  |  |
| E7 | 1.87e-08 | 9.49e-30 | 3.66e-60 | 9.84e-43 | 1.76e-09 | 2.14e-22 |  |
| E8 | 3.55e-28 | 3.96e-59 | 3.97e-103 | 1.39e-78 | 1.11e-31 | 1.91e-50 | 1.40e-08 |

**Table S8: variability (distance to median) comparison between any two stages**

Wilcoxon test, multiple test corrected *p*-values (Benjamini–Hochberg method).

| Stages | E1 | E2 | E3 | E4 | E5 | E6 | E7 |
| --- | --- | --- | --- | --- | --- | --- | --- |
| E2 | 5.64e-51 |  |  |  |  |  |  |
| E3 | 4.83e-94 | 3.00e-04 |  |  |  |  |  |
| E4 | 3.57e-54 | 1.13e-01 | 3.90e-10 |  |  |  |  |
| E5 | 9.54e-01 | 2.03e-61 | 1.09e-119 | 4.91e-70 |  |  |  |
| E6 | 1.58e-11 | 1.82e-18 | 9.34e-52 | 3.13e-24 | 2.71e-11 |  |  |
| E7 | 2.41e-30 | 1.35e-154 | 5.48e-265 | 5.28e-203 | 3.12e-44 | 5.15e-87 |  |
| E8 | 9.09e-98 | 1.10e-251 | 0.00e+00 | 0.00e+00 | 9.77e-148 | 2.36e-210 | 1.22e-37 |

**Table S9: bootstrapped median variability comparison between any two stages**

Wilcoxon test, multiple test corrected *p*-values (Benjamini–Hochberg method).

| Stages | E1 | E2 | E3 | E4 | E5 | E6 | E7 |
| --- | --- | --- | --- | --- | --- | --- | --- |
| E2 | 1.60e-66 |  |  |  |  |  |  |
| E3 | 7.73e-99 | 3.94e-22 |  |  |  |  |  |
| E4 | 2.26e-96 | 1.38e-18 | 2.41e-18 |  |  |  |  |
| E5 | 1.14e-02 | 4.67e-134 | 1.20e-303 | 4.16e-163 |  |  |  |
| E6 | 1.50e-40 | 1.33e-27 | 2.62e-112 | 1.17e-111 | 3.11e-123 |  |  |
| E7 | 4.58e-90 | 2.51e-163 | 4.15e+00 | 4.15e-164 | 2.96e-141 | 1.05e-163 |  |
| E8 | 4.15e-139 | 1.05e-163 | 4.15e+00 | 4.15e-164 | 7.64e-159 | 1.20e-163 | 2.60e-59 |

**Table S10: variability (adjusted SD) comparison between any two stages (Genes with constant expression level over development)**

Wilcoxon test, multiple test corrected *p*-values (Benjamini–Hochberg method).

| Stages | E1 | E2 | E3 | E4 | E5 | E6 | E7 |
| --- | --- | --- | --- | --- | --- | --- | --- |
| E2 | 2.19e-09 |  |  |  |  |  |  |
| E3 | 3.00e-07 | 2.98e-01 |  |  |  |  |  |
| E4 | 5.26e-08 | 7.12e-01 | 5.93e-01 |  |  |  |  |
| E5 | 2.19e-01 | 8.97e-08 | 7.50e-06 | 4.15e-06 |  |  |  |
| E6 | 4.09e-02 | 1.21e-05 | 3.74e-04 | 2.09e-04 | 4.56e-01 |  |  |
| E7 | 7.19e-11 | 1.47e+43 | 1.55e+39 | 1.49e-37 | 1.16e-18 | 5.29e-21 |  |
| E8 | 7.66e-17 | 5.29e+47 | 1.47e+43 | 8.04e-43 | 1.57e-26 | 8.25e-29 | 7.00e-04 |

**Table S11: variability (adjusted SD) comparison between any two stages (transcription factors)**

Wilcoxon test, multiple test corrected *p*-values (Benjamini–Hochberg method).

| Stages | E1 | E2 | E3 | E4 | E5 | E6 | E7 |
| --- | --- | --- | --- | --- | --- | --- | --- |
| E2 | 6.44e-29 |  |  |  |  |  |  |
| E3 | 3.16e-35 | 2.02e-01 |  |  |  |  |  |
| E4 | 5.40e-31 | 5.62e-01 | 5.06e-01 |  |  |  |  |
| E5 | 1.88e-14 | 1.73e-06 | 1.76e-09 | 3.29e-07 |  |  |  |
| E6 | 3.66e-34 | 2.63e-01 | 9.15e-01 | 5.62e-01 | 1.21e-08 |  |  |
| E7 | 2.40e-02 | 2.58e-56 | 6.63e-65 | 4.55e-58 | 1.87e-33 | 4.07e-62 |  |
| E8 | 9.00e-19 | 2.55e-94 | 1.19e-104 | 2.79e-97 | 7.45e-74 | 5.73e-103 | 2.46e-17 |

**Table S12: variability (adjusted SD) comparison between any two stages (dispersed promoter genes)**

Wilcoxon test, multiple test corrected *p*-values (Benjamini–Hochberg method).

| Stages | E1 | E2 | E3 | E4 | E5 | E6 | E7 |
| --- | --- | --- | --- | --- | --- | --- | --- |
| E2 | 1.04e-174 |  |  |  |  |  |  |
| E3 | 8.26e-198 | 3.36e-01 |  |  |  |  |  |
| E4 | 1.49e-196 | 3.44e-02 | 2.65e-01 |  |  |  |  |
| E5 | 2.60e-36 | 3.19e-75 | 3.63e-90 | 1.22e-89 |  |  |  |
| E6 | 1.07e-124 | 3.55e-07 | 1.08e-10 | 2.09e-12 | 1.85e-37 |  |  |
| E7 | 1.18e-41 | 0.00e+00 | 0.00e+00 | 0.00e+00 | 8.44e-189 | 0.00e+00 |  |
| E8 | 4.79e-168 | 0.00e+00 | 0.00e+00 | 0.00e+00 | 0.00e+00 | 0.00e+00 | 1.61e-82 |

**Table S13: variability (adjusted SD) comparison between any two stages (precise promoter genes)**

Wilcoxon test, multiple test corrected *p*-values (Benjamini–Hochberg method).

| Stages | E1 | E2 | E3 | E4 | E5 | E6 | E7 |
| --- | --- | --- | --- | --- | --- | --- | --- |
| E2 | 5.31e-74 |  |  |  |  |  |  |
| E3 | 2.34e-183 | 2.64e-33 |  |  |  |  |  |
| E4 | 1.81e-94 | 1.13e-02 | 3.07e-21 |  |  |  |  |
| E5 | 3.79e-01 | 3.42e-93 | 4.74e-218 | 1.60e-115 |  |  |  |
| E6 | 1.90e-30 | 1.08e-12 | 2.26e-79 | 3.49e-22 | 4.61e-39 |  |  |
| E7 | 4.74e-24 | 1.35e-213 | 0.00e+00 | 3.61e-248 | 2.58e-24 | 8.48e-121 |  |
| E8 | 2.20e-116 | 0.00e+00 | 0.00e+00 | 0.00e+00 | 2.40e-123 | 5.79e-265 | 2.56e-67 |

**Table S14: proximal promoter H3K4Me1 signal (Z score) comparison between any two stages**

Wilcoxon test, multiple test corrected *p*-values (Benjamini–Hochberg method).

| Satges | 0-4h | 4-8h | 8-12h | 12-16h | 16-20h |
| --- | --- | --- | --- | --- | --- |
| 4-8h | 2.31e-63 |  |  |  |  |
| 8-12h | 2.20e-72 | 4.13e-191 |  |  |  |
| 12-16h | 8.34e-24 | 5.35e-06 | 9.00e-155 |  |  |
| 16-20h | 1.94e-104 | 2.90e-248 | 3.68e-02 | 2.73e-175 |  |
| 20-24h | 2.08e-146 | 3.01e-01 | 1.98e-228 | 6.96e-29 | 0.00e+00 |

**Table S15: gene body H3K4Me1 signal (Z score) comparison between any two stages**

| Satges | 0-4h | 4-8h | 8-12h | 12-16h | 16-20h |
| --- | --- | --- | --- | --- | --- |
| 4-8h | 2.31e-210 |  |  |  |  |
| 8-12h | 2.73e-20 | 1.68e-285 |  |  |  |
| 12-16h | 1.68e-114 | 4.53e-06 | 1.99e-198 |  |  |
| 16-20h | 3.20e-158 | 0.00e+00 | 9.19e-25 | 0.00e+00 |  |
| 20-24h | 1.01e-33 | 6.70e-127 | 4.66e-65 | 9.74e-45 | 0.00e+00 |

**Table S16: proximal promoter H3K27Ac signal (Z score) comparison between any two stages**

Wilcoxon test, multiple test corrected *p*-values (Benjamini–Hochberg method).

| Satges | 0-4h | 4-8h | 8-12h | 12-16h | 16-20h |
| --- | --- | --- | --- | --- | --- |
| 4-8h | 2.45e-03 |  |  |  |  |
| 8-12h | 7.91e-141 | 2.12e-88 |  |  |  |
| 12-16h | 6.90e-01 | 1.85e-04 | 4.74e-127 |  |  |
| 16-20h | 8.98e-01 | 3.97e-03 | 6.20e-147 | 6.87e-01 |  |
| 20-24h | 2.97e-103 | 8.33e-85 | 6.08e-292 | 1.31e-57 | 1.89e-165 |

**Table S17: gene body H3K27Ac signal (Z score) comparison between any two stages**

| Satges | 0-4h | 4-8h | 8-12h | 12-16h | 16-20h |
| --- | --- | --- | --- | --- | --- |
| 4-8h | 1.05e-22 |  |  |  |  |
| 8-12h | 3.37e-74 | 3.79e-129 |  |  |  |
| 12-16h | 4.25e-25 | 3.90e-02 | 5.99e-142 |  |  |
| 16-20h | 1.02e-27 | 4.42e-79 | 3.52e-33 | 4.82e-76 |  |
| 20-24h | 7.30e-27 | 8.55e-03 | 3.57e-154 | 7.48e-03 | 3.85e-177 |

**Table S18: proximal promoter H3K9Ac signal (Z score) comparison between any two stages**

Wilcoxon test, multiple test corrected *p*-values (Benjamini–Hochberg method).

| Satges | 0-4h | 4-8h | 8-12h | 12-16h | 16-20h |
| --- | --- | --- | --- | --- | --- |
| 4-8h | 1.73e-06 |  |  |  |  |
| 8-12h | 2.17e-40 | 3.75e-16 |  |  |  |
| 12-16h | 9.49e-01 | 1.71e-07 | 2.17e-40 |  |  |
| 16-20h | 4.50e-78 | 2.03e-97 | 1.23e-171 | 1.41e-67 |  |
| 20-24h | 7.31e-145 | 3.75e-148 | 1.11e-230 | 8.16e-121 | 1.13e-28 |

**Table S19: gene body H3K9Ac signal (Z score) comparison between any two stages**

| Satges | 0-4h | 4-8h | 8-12h | 12-16h | 16-20h |
| --- | --- | --- | --- | --- | --- |
| 4-8h | 3.11e-12 |  |  |  |  |
| 8-12h | 2.59e-11 | 1.03e-34 |  |  |  |
| 12-16h | 4.27e-40 | 1.40e-12 | 7.11e-77 |  |  |
| 16-20h | 2.65e-10 | 2.22e-01 | 1.28e-38 | 2.94e-15 |  |
| 20-24h | 2.30e-21 | 3.02e-01 | 1.52e-53 | 2.77e-08 | 1.23e-05 |
